# Automated, Reproducible Investigation of gene set Differential Enrichment via the AUTO-go framework

**DOI:** 10.1101/2022.02.25.482003

**Authors:** Eleonora Sperandio, Isabella Grassucci, Lorenzo D’Ambrosio, Matteo Pallocca

**Affiliations:** UOSD Biostatistics, Bioinformatics and Clinical Trial Center, IRCSS Regina Elena National Cancer Institute, Rome; UOSD Immunology and Tumor Immunotherapy, IRCSS Regina Elena National Cancer Institute, Rome

## Abstract

Reproducibility in Life Sciences is challenged in the analysis of large multi-omics datasets. One of the final steps of said processes is Gene Set enrichment, where web tools represent a valuable resource but not a reliable surrogate for standardized, high-quality visualizations. The AUTO-go framework proposes standardization of the Gene Functional Enrichment process along with an R framework able to produce high-quality visualization in an automated manner, improving the reproducibility of the whole analytical process. We present three use cases in Cancer Transcriptomics and Epigenomics datasets as a proof-of-concept to visualize Multiple Differential Expression and Single Sample Gene Set Enrichment Analysis.

**Author Summary:** Bioinformatics and Data Science are routinely challenged to distill intelligible results from huge amounts of data. These results, in turn, are conveyed through plots and visualizations that should be easily reproducible for scientific soundness and ethical reasons. A specific area in which these analyses are of critical importance is Genomics, where Genes functions need to be enriched when comparing pathological states or treatments. Here we present a software framework that aims at standardizing said differential analyses and visualizations when dealing with genomics data. Finally, we show how it can be employed to shear light on publicly available datasets, even in small casuistry of Rare Cancers.

## Introduction

Gene Ontology (GO) and Pathway Enrichment Analysis are pivotal aspects of Life Science research – but the level of standardization and reproducibility is worryingly low for such popular techniques [1].

Additionally, most of the enrichment analyses currently published rely on web applications that, on the one hand, enable non-bioinformaticians to conduct exploratory analyses; on another, open concern for result reproducibility, being a *manual* step of data processing strongly contrasting the rules for reproducible bioinformatics [2-3].

Virtualization techniques such as Docker and Singularity helped to encapsulate software enabling total reproducibility, while additional workflow management layers such as Nextflow and Snakemake [4-5] enabled to build of complex virtualized pipelines and run them in HighPerformance Computing Clusters. Unfortunately, what is presented on a life science paper is not primary output matrices, but functional enrichments that currently do not benefit from such advancements.

Among the R packages available to the community, the *clusterProfiler* is a notable exception, with a development that has focused many features on genomics coordinates enrichment and specific high-throughput experiments, while our focus lies on the high-level conceptualization and visualization of differential analysis [6].

Here we present AUTO-go, a logical and bioinformatics framework that enables (1) reproducible GO analyses; (2) high quality automated visualizations; (3) proposes a high-level visualization for complex experimental designs with multiple comparisons.

### Design and implementation

According to the logical framework (Fig. 1), a Differential Expression is the most frequent starting input from which one or several gene lists are extracted according to fold change and statistical significance filters (e.g., *strongly upregulated, log2FC > 1 and padj < 0*.*05)*. The protocol core is an atomic function that enriches a gene list over a list of selected databases, from which several visualizations are produced (Fig 1). The gene list can derive from several Genomics applications as described in the Use Cases section.

**Fig 1.**
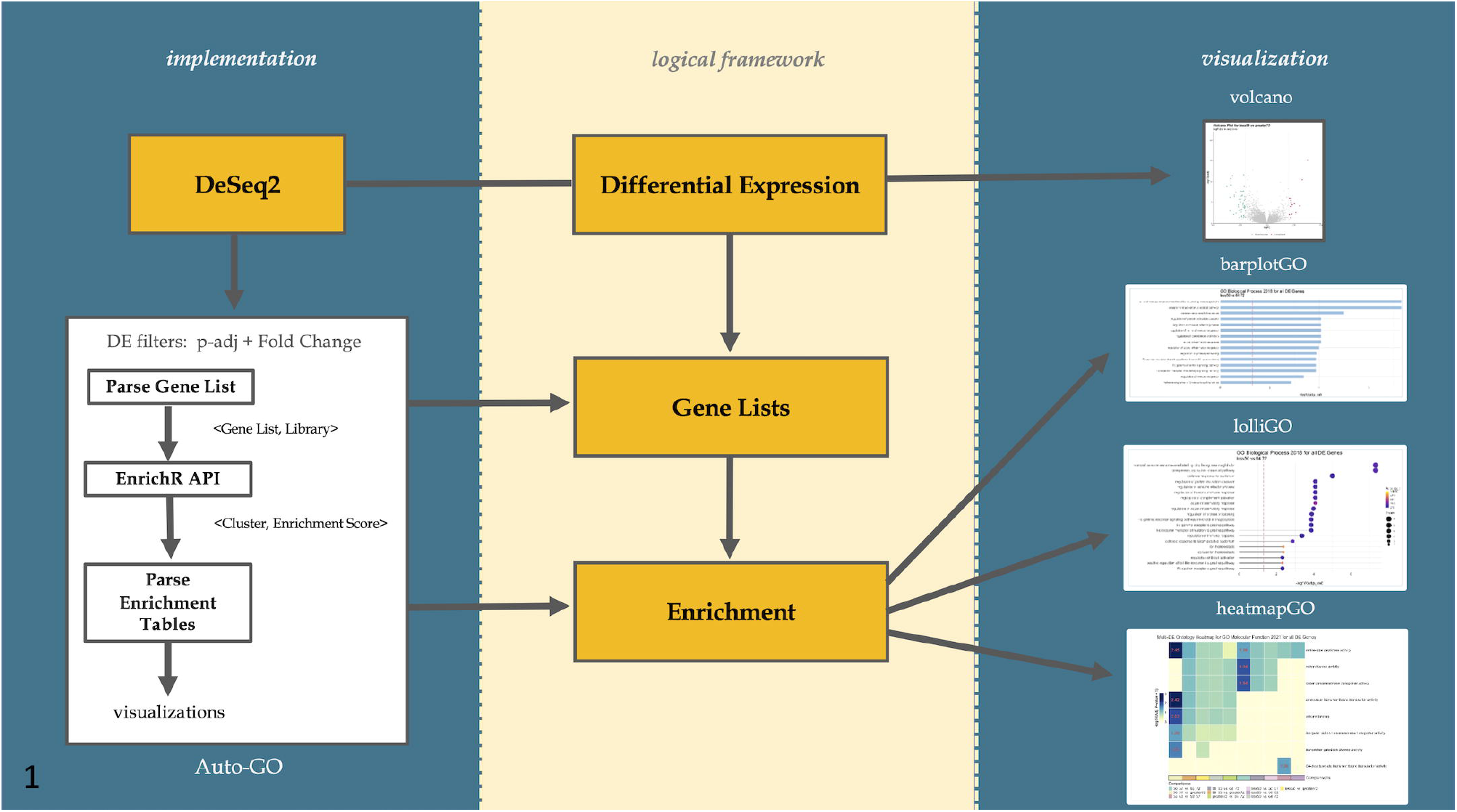
Logical framework and implementation workflow.

### Gene List Enrichment Visualization

Every *<gene list, database>* combination produces a high-quality bar plot with the top *N* terms enriched, with dynamic resizing to accommodate long terms naming in the final plot.

The current implementation of the core module relies on the Enrichr API [7], but it is engineered to be generalized with other enrichment functions, with the only constrain of having a gene list as input and a tuple matrix with *<Term, Enrichment Score>* as output.

### Multiple Comparison Visualization

A classical need in -omics analysis is the representation of functional terms enriched in several conditions or comparisons. The HeatmapGO module is built to provide a high-level visualization of multiple comparisons enrichment, with rows representing terms, such as GO components and Transcription Factors, and columns being experimental comparisons.

### ssGSEA

In many fields, the scarcity of sample availability does not allow classical statistical modeling. The challenge in obtaining robust results is exacerbated by the employment of -omics profiling, collecting thousands of features per observation. To this purpose, we expanded the AUTO-go package with the single-sample implementation of the Gene Set Enrichment Analysis Algorithm [8-9], allowing researchers to compare discrete cohorts of samples over known gene signatures.

For all the visualization depicting a subset of the enriched terms, a ranking choice must be made to represent a human-readable number of terms and clusters. In the ssGSEA and HeatmapGO, the top 20 terms are selected by ascending -log10(p-adjusted) score. Other developers and data scientists would pick a different ranking employing a mixture of significance and variance among samples and comparisons to show the functions having a strong modulation.

## Results

To provide a proof-of-concept application of our package, we envisioned three analytical settings to test it, namely 1. Large RNA-seq casuistry with multiple comparisons Differential Expression and Enrichment (Tumor Cancer Genome Atlas, TCGA) 2. Discrete in-vitro enrichment of gene lists representing epigenetic signals (Encyclopedia of DNA elements, ENCODE) 3. Discrete in-vivo rare tumor samples profiled via total RNA-seq. (Fig. 2). In the first use case, the TCGA dataset of Skin Cutaneous Melanoma Adenocarcinoma (TCGA-SKCM) was partitioned according to a specific immunotherapy biomarker, the Tumor Mutational Burden (TMB). Differential expression was carried out by comparing all TMB quartiles (Fig. 2,3) [10]. The KEGG 2021 Heatmap shows a stronger enrichment in Ras signaling pathway in the higher group comparison (Q3-Q4), suggesting a switch in the higher mutational load group (Fig. 2,3). This enrichment can be further investigated at the LolliGO level showing that most Ras-related genes are upregulated except for FGF5, while down-regulated genes are more enriched in Cortisol synthesis and secretion, less evident from the Heatmap.

**Fig 2.**
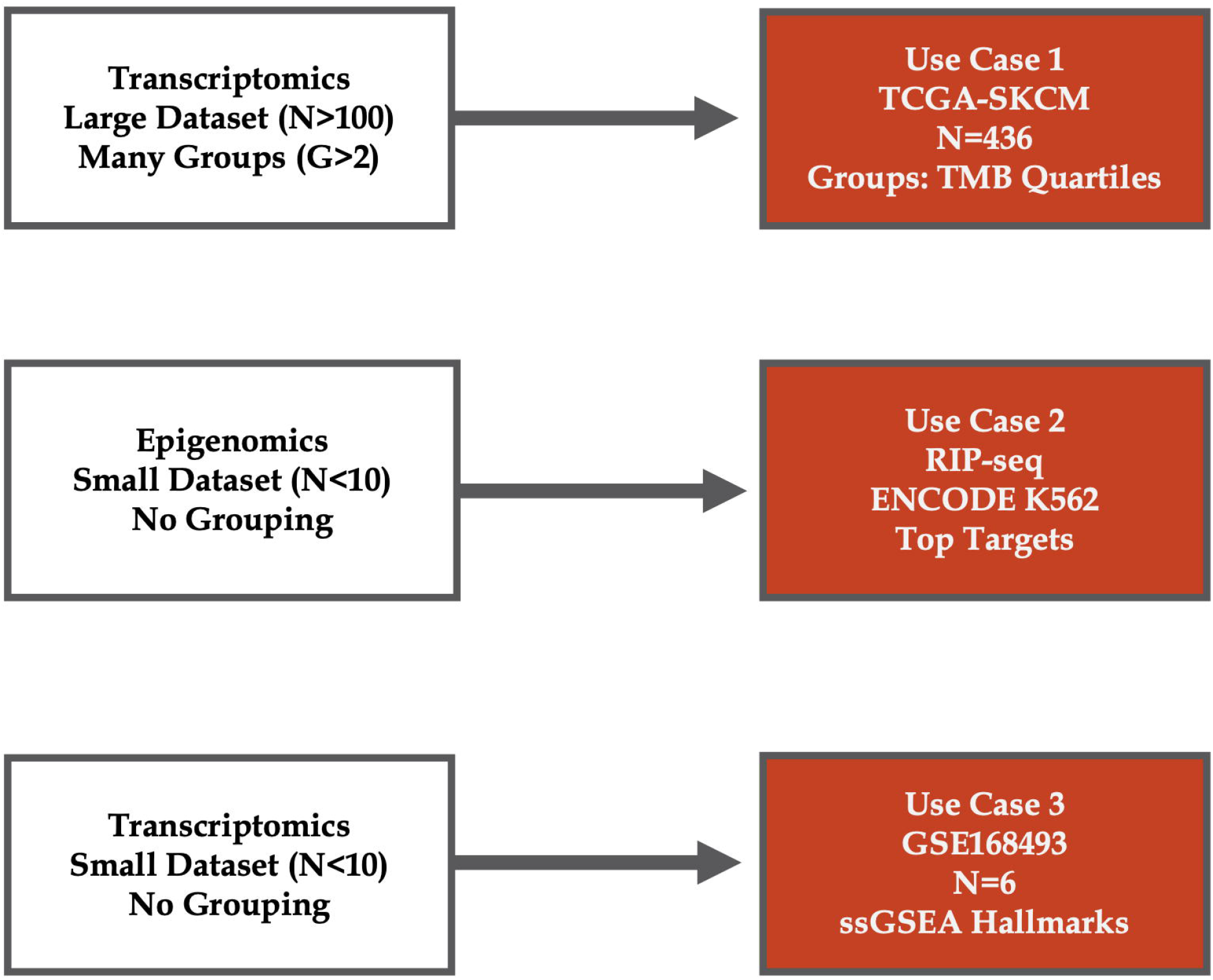
Use case schema. Workflow of the three use cases with different casuistry and comparison sizes.

**Fig 3.**
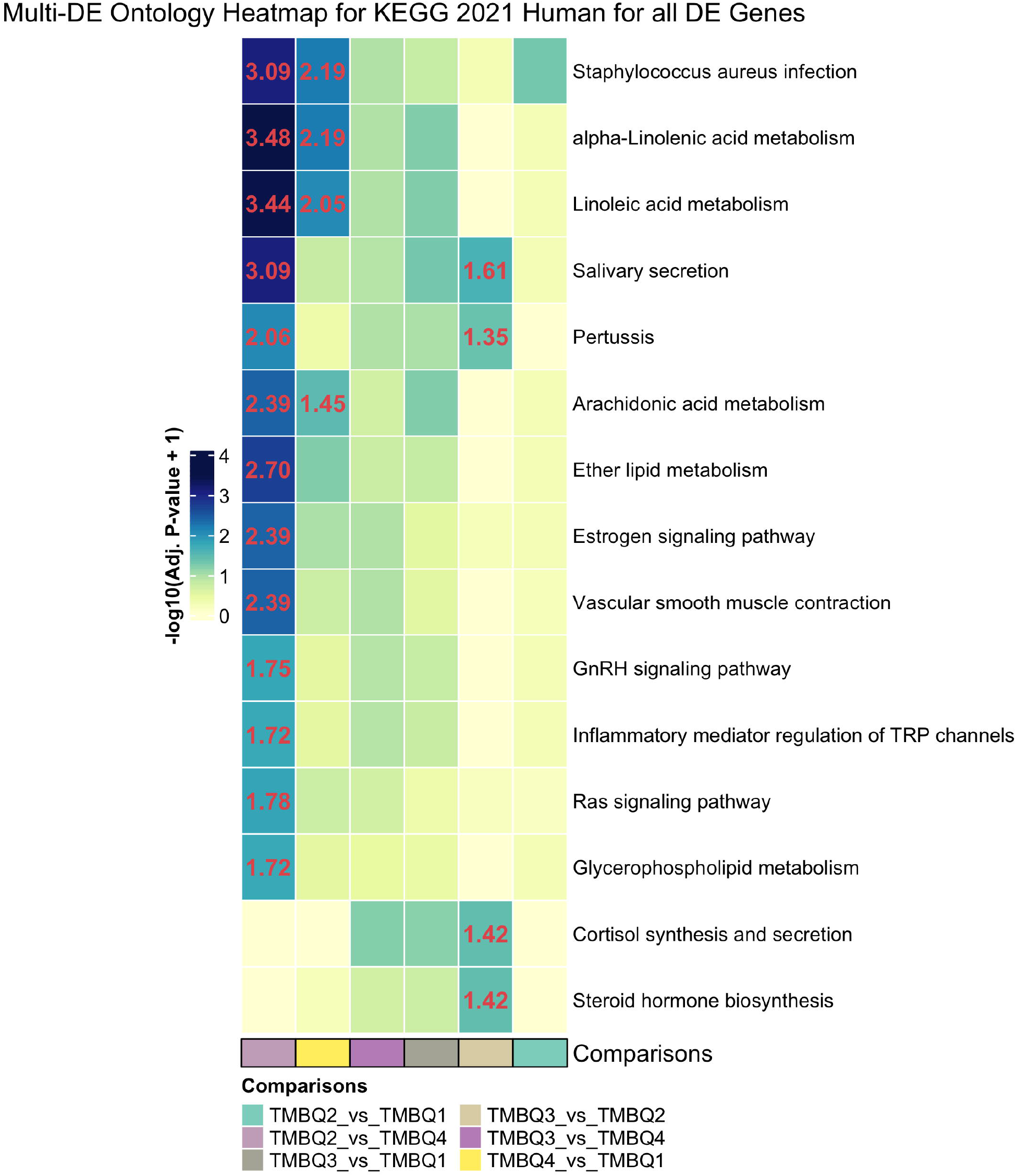
Results on DE genes enriched on the TCGA-SKCM multiple TMB comparisons. Rows: enriched terms over KEGG_2021 Enrichr library. Columns: genes regulated in multiple comparisons. Cell content: *-log10(padj + 1)* reported only for significant clusters.

Next, the epigenetic unit test was fetched from the ENCODE database, fetching all the RNA Immunoprecipitation sequencing (RIP-seq) available in the K562 cell line. Gene lists were obtained by annotating with Homer [11] the enriched peaks and extracting only the promoter-TSS records. In this scenario, the Cellular Component database coupled with the *lolliGO* modules shows stronger enrichment hydrogen peroxide metabolic and catabolic process in mRNA targets of ELAVL1 in (Fig. 4).

**Fig 4.**
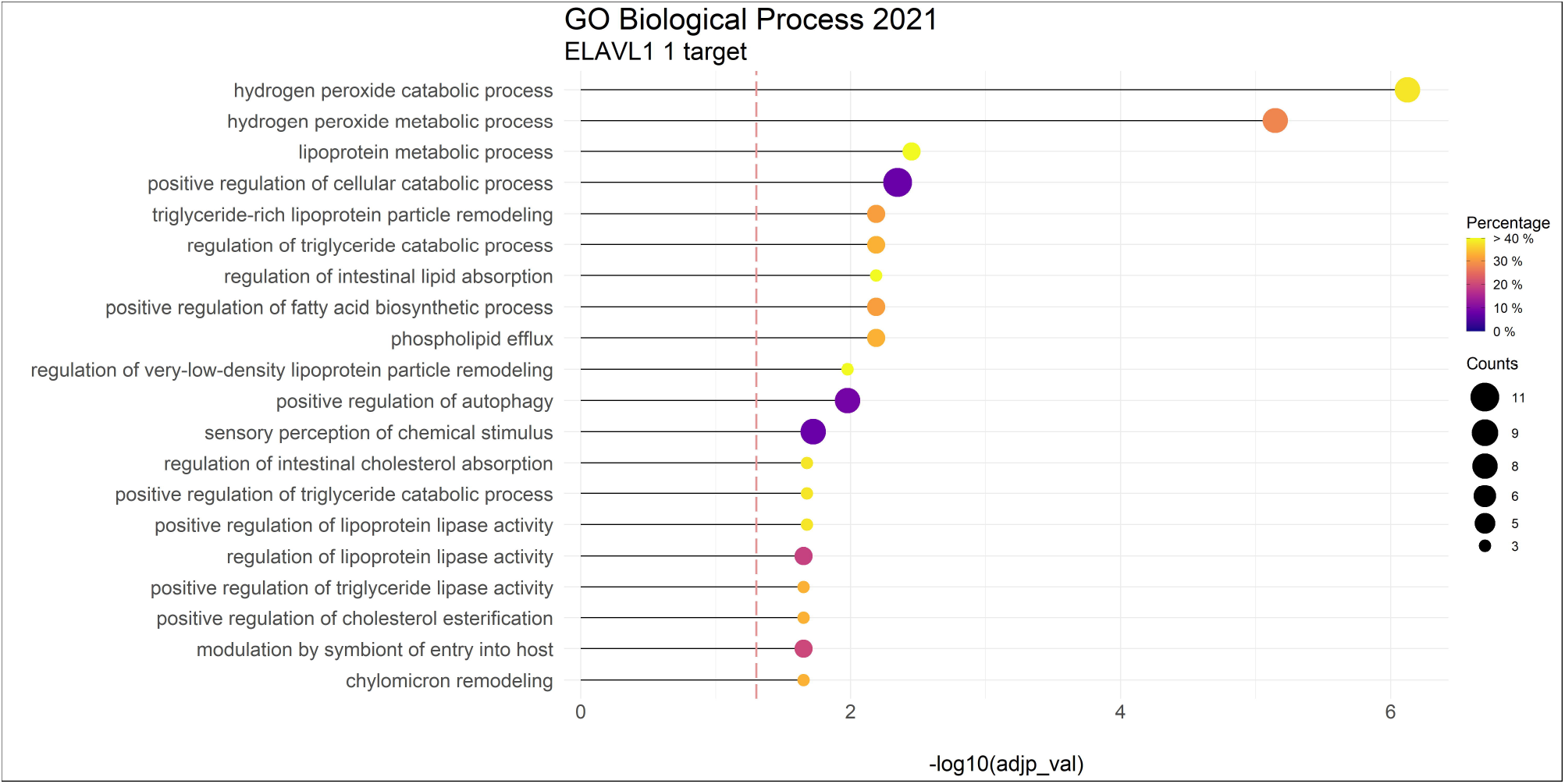
LolliGO plot from RIP-seq: Ontology enrichment over a list of ELAVL targets derived from RIP-seq. Color: percentage of the cluster given as input with respect to the total functional cluster. Dot size: gene count for cluster.

Finally, the third unit test was carried out on the on the GSE168493 record, containing total RNA-seq profiles from a small casuistry of Epithelioid hemangioendothelioma, a rare tumor with an incidence of 1 out of million people [12]. In this instance, the ssGSEA package enables to shear light into the pathway activation peculiarities of said tumors, with a stronger enrichment of PSMB5 target genes in samples hEHE.6 and hEHE5 (Fig. 5).

**Fig 5.**
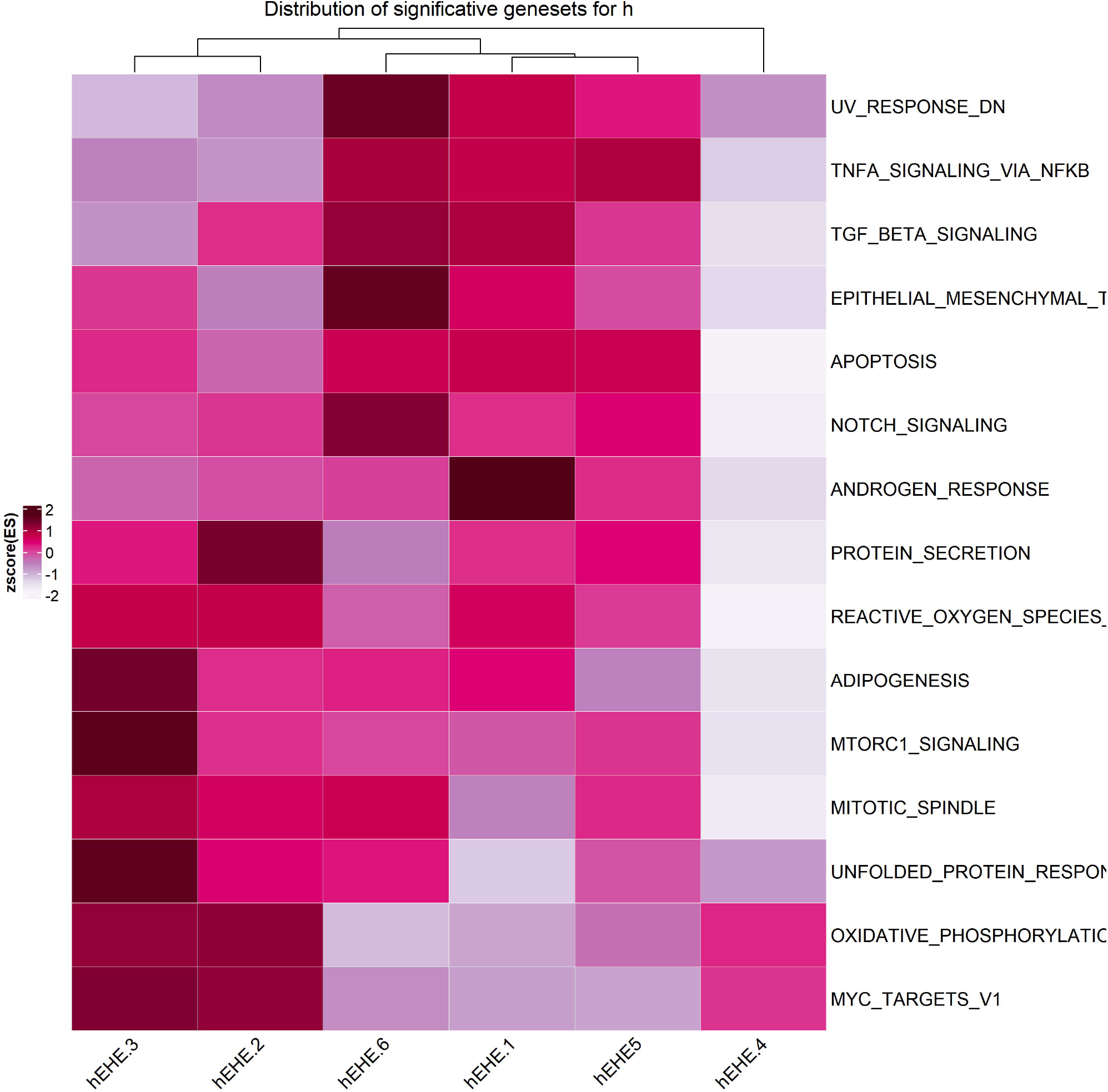
Single-Sample Gene Set Enrichment Analysis heatmap: Heatmap showing ssGSEA enrichment over the *Hallmark* term for the 6 RNA-seq samples (eEHE1-6). Z-score of the Enrichment Score in cell content.

Taken together, all these examples point out many analytical scenarios in which the Auto-GO package can provide a solid foundation and a valuable engineering tool for -omics-focused Bioinformaticians.

## Availability and Future Directions

The package is available at https://gitlab.com/bioinfo-ire-release/auto-go. The repository contains a step-by-step tutorial for the whole framework usage and the data input to reproduce the first use case presented in the results section, along with a Dockerfile. All the generated outputs, folders, and figures are available in the tutorial and in Fig. S1.

**Fig. S1.**
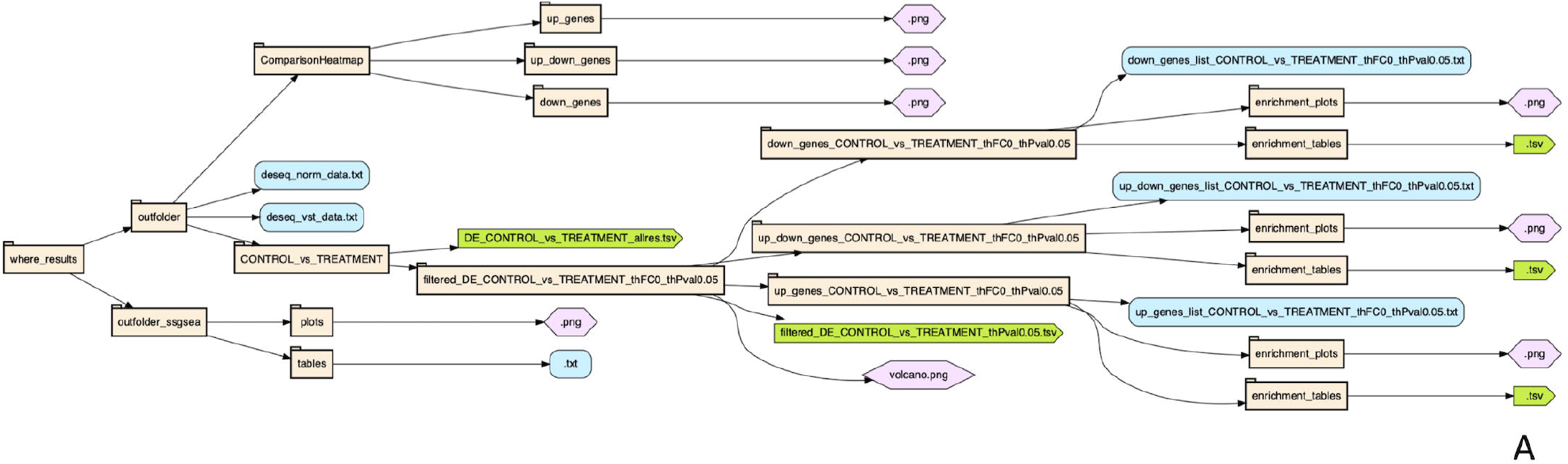
Folder tree of the AUTO-go output.

## Acknowledgements

We thank Francesca Nardozza for editorial assistance.

## Author Contributions

Conceptualization: Matteo Pallocca, Eleonora Sperandio

Formal analysis: Matteo Pallocca, Eleonora Sperandio

Methodology: Matteo Pallocca, Eleonora Sperandio, Isabella Grassucci, Lorenzo D’Ambrosio

Software: Eleonora Sperandio

Supervision: Matteo Pallocca

Validation: Lorenzo D’Ambrosio, Isabella Grassucci

Writing – original draft: Matteo Pallocca

Writing – review & editing: Matteo Pallocca, Isabella Grassucci

## Funding

This work has been supported by the Italian Ministry of Health (Ricerca Corrente 2020).

## Conflict of Interest

none declared.

## Notes

https://gitlab.com/bioinfo-ire-release/auto-go

### Competing Interest Statement

The authors have declared no competing interest.

